# Vinculin plays a role in neutrophil stiffening and transit through model capillary segments

**DOI:** 10.1101/2022.04.24.489286

**Authors:** Brittany M. Neumann, Zachary S. Wilson, Kinga Auguste, Yasmin Roye, Manisha K. Shah, Eric M. Darling, Craig T. Lefort

**Affiliations:** Division of Surgical Research, Department of Surgery, Rhode Island Hospital, Providence, RI 02903, USA; Graduate Program in Pathobiology, Brown University, Providence, RI 02912, USA; Center for Biomedical Engineering, Brown University, Providence, RI 02912, USA; Department of Pathology and Laboratory Medicine, Department of Orthopaedics, School of Engineering, Brown University, Providence, RI 02912, USA

**Keywords:** Neutrophils, Cell signaling, Actin cytoskeleton, Deformability, Microfluidics

## Abstract

Neutrophils are rapidly mobilized from the circulation to sites of inflammation. The mechanisms of neutrophil trafficking in the lung are distinct from those in the periphery, in part because the pulmonary capillaries are the primary site of neutrophil emigration rather than postcapillary venules. Since the diameter of a neutrophil is greater than the width of most pulmonary capillary segments, they must deform to transit through this capillary network, even at homeostasis. Resistance to deformation is primarily due to cortical actin that is rapidly assembled when a neutrophil is exposed to a priming or activation stimulus, resulting in neutrophil stiffening and subsequent sequestration within the pulmonary capillary network. In the current study, we use a microfluidic assay to characterize neutrophil transit through model capillary-like channels. Using techniques from single-particle tracking, we analyzed the cumulative distribution of neutrophil transit times and resolve population-based effects. We found that vinculin, an actin-binding adaptor protein, plays an essential role in neutrophil stiffening in response to formyl-Met-Leu-Phe (fMLP). Vinculin-deficient neutrophils lack the development of a population with slow transit through narrow channels that was observed in both wild-type murine bone marrow neutrophils and HoxB8-conditional progenitor-derived neutrophils. Atomic force microscopy studies provide further evidence that vinculin is required for neutrophil stiffening. Consistent with these findings, we observed that neutrophil sequestration in the lungs of mice is attenuated in the absence of vinculin. Together, our studies indicate that vinculin mediates actin-dependent neutrophil stiffening that leads to their sequestration in capillaries.

## Introduction

Acute respiratory distress syndrome (ARDS) has an in-hospital mortality rate of up to 40%, and few effective pharmacological therapies exist [1-3]. Part of the difficulty with ARDS treatment is due to its heterogeneity that arises from a wide range of triggers, its broad clinical definition, and variable outcomes [4]. The pathophysiological hallmarks include severe inflammatory injury to the alveolar-capillary barrier, surfactant depletion, loss of aerated lung tissue, and neutrophil infiltration into the lung [5, 6]. Long-term mortality ranges from 11% to 60%, and survivors often have significantly reduced quality of life with sequelae including diffuse pulmonary fibrosis, chronic lung disease, neuropsychiatric impairments, critical illness associated polyneuropathy, myopathy, and neuromyopathy [7, 8].

There is abundant evidence for the pathophysiological role of dysregulated neutrophil trafficking and activation in ARDS [9]. Activated neutrophils release granular enzymes, reactive oxygen species (ROS), neutrophil extracellular traps (NETs), and pro-inflammatory cytokines in response to stimuli. In an ideal host response, neutrophils deploy an antimicrobial armament once it reaches the site of infection. However, when this response becomes dysregulated, neutrophils can accumulate in the lungs and indiscriminately release cytotoxic species, leading to acute lung injury (ALI) and potentially ARDS [5, 6].

Circulating neutrophils passing through the lung exhibit unique capture and trafficking characteristics due to the geometrical features of the microvasculature [10-12]. Capillaries envelop each alveolus and have diameters of 2-15 μm, which is frequently less than the 8-11 μm diameter of a neutrophil [13-15]. Neutrophils must repeatedly deform to pass through the pulmonary capillaries, slowing their transit relative to red blood cells and resulting in their enrichment within the vascular bed of the lungs [10, 11]. These geometrical characteristics of the lung microvasculature result in altered neutrophil homing mechanisms compared to those in the periphery. In contrast to selectin-mediated neutrophil rolling that occurs in postcapillary venules, the initial capture and enrichment of neutrophils in the pulmonary system is due to these cells’ mechanical properties (i.e., stiffness) [10, 16]. Additionally, the prominence and role of β2 integrins in neutrophil interactions within the lung vary depending on inflammation conditions [12, 17-19].

Neutrophil stiffness arises from four cellular components: the plasma membrane, which has a low shear modulus and is elastic but also antagonizes actin-based protrusions [20]; the actin cortex, which provides cortical tension and cellular structure; the cytoplasm, which has viscous resistance; and the nucleus, which is modeled as an immiscible Newtonian liquid drop [13]. Each of these components plays a different role in defining the elastic modulus of the cell. The plasma membrane and the interacting actin cortex are the leading players in neutrophil deformation under the time scales and pressures we are investigating.

Upon exposure to stimuli, neutrophils reorganize their actin cytoskeleton and polymerize F-actin [21, 22]. These actin-dependent changes reduce neutrophil deformability, resulting in sequestration and prolonged retention within the pulmonary capillaries [10, 23-26]. Although there is abundant evidence linking cortical actin development to neutrophil stiffness, much remains unknown about the biochemical mechanisms responsible for organizing this actin-dependent phenotype. Vinculin is a non-enzymatic scaffolding protein that binds to the actin cytskeleton and is involved in its reorganization during dynamic processes such as cell migration [27]. In this study, we describe the role of vinculin in mediating neutrophil stiffening, prolonging their transit through microfluidic constrictions that model the pulmonary capillary unit.

## Materials and Methods

### Antibodies and reagents

All antibodies used are against murine antigens. Antibodies: PE-anti-CD117 (clone ACK2; BioLegend), APC-anti-Ly6G (clone 1A8; BioLegend), anti-vinculin (Cell Signaling Technologies), HRP-conjugated-anti-Rabbit IgG (Cell Signaling Technologies). Reagents: Zombie Violet Fixable Viability Kit (BioLegend), recombinant murine SCF (BioLegend), recombinant murine G-CSF (BioLegend), N-Formyl-Met-Leu-Phe (fMLP), BioXtra ≥ 99.0% (Millipore Sigma), 4-Hydroxytamoxifen (Tocris), carboxyfluorescein succinimidyl ester (CFSE) (BioLegend), SYLGARD 184 Silicone Elastomer Kit, polydimethylsiloxane (PDMS) (Dow Corning), pHrodo Green *S. aureus* Bioparticles™ Conjugate for Phagocytosis (Invitrogen), Gibco Opti-MEM reduced serum media (ThermoFisher Scientific), Fetal Bovine Serum (FBS) — heat inactivated (GeminiBio), 2-mercaptoethanol (Millipore Sigma), penicillin-streptomycin (ThermoFisher Scientific), MEM non-essential amino acids (NEAA) (ThermoFisher Scientific), phosphate-buffered saline (PBS) (ThermoFisher Scientific), Hanks’ Balanced Salt Solution containing Ca^2+^/Mg^2+^ (HBSS^++^) (ThermoFisher Scientific), Pluronic F-127 (Millipore Sigma), bovine serum albumin (BSA) lyophilized powder ≥ 96.0% (Millipore Sigma), DNAaseI (Zymo Research), trichloro(1H, 1H, 2H, 2H-perfluorooctyl)silane (Millipore Sigma), blasticidin (Tocris), puromycin (Tocris).

### HoxB8-conditional murine neutrophil progenitors

In brief, murine hematopoietic progenitors were isolated from bone marrow with the EasySep Mouse HSPC Enrichment Kit (StemCell Technologies) and transduced with a tamoxifen-inducible expression vector *HoxB8*. The basal media conditions used for cell culture were as follows: Opti-MEM, 10% FBS, NEAA, pen/strep, and 30 μM 2-mercaptoethanol. To maintain cells in the progenitor state, cells were cultured in the presence of 100 nM 4-hydroxytamoxifen (4-OHT), 50 ng/mL recombinant murine stem cell factor (mSCF), and 1 μg/mL puromycin for selection of the transduced cells. The surviving cells made up our WT progenitor cell lines [28-30]. In some cases, 2% conditioned media from CHO cells producing SCF was used in place of mSCF (a gift from Dr. Patrice Dubreuil).

Differentiation of progenitors into neutrophils was performed by washing the cells extensively with PBS and resuspending them in basal media containing 20 ng/mL mSCF and 20 ng/mL recombinant murine granulocyte colony-stimulating factor (mG-CSF) for two days. On day two, cells were washed and resuspended in basal media with 20 ng/mL mG-CSF for three days. Successfully differentiated neutrophils exhibit multi-lobed nuclei, expression of Ly6G, and loss of expression of CD117 (cKit) [28-30].

### Gene disruption

To generate a vinculin knockout (Vcl^-/-^) progenitor cell line, HoxB8-conditional progenitors were transduced with a lentiviral vector that expressed Cas9 and single-guide RNA (sgRNA) targeting the *Vcl* gene. We modified the pLentiCRISPR v2 vector (gift from Feng Zhang – Addgene plasmid #52961) to confer blasticidin resistance and the following sgRNA target sequences: CCGGCGCGCTCACCCGGACG [29]. Empty vector expression of Cas9 without targeting sgRNA was used as wild-type (WT) control. Vcl^-/-^ was selected via blasticidin selection at 5 μg/mL concentration. Gene disruption was confirmed via western blot [29].

### Atomic Force Microscopy (AFM)

AFM was performed using similar methods to those previously published [31, 32]. Coverslips (#1.5) were plasma cleaned before adding neutrophils. *In vitro*-derived murine neutrophils were differentiated and used on day 4. Neutrophils were stimulated for 15 minutes in HBSS^++^ with either vehicle control or 1 µM fMLP at 37°C.

Cantilevers were purchased from Novascan Technologies with 5 µm borosilicate beads (k ∼ 0.03 N/m). Cantilever spring constants were determined by the power spectral density of thermal noise fluctuations. Cantilevers were used to probe neutrophils using elastic indentation tests with an approach velocity of 10 µm/s. A 0.6 nN trigger force was used to limit indentations to less than 10% strain based on the cell’s height. All tests were done at room temperature. A modified Hertz model (Equation 1) was used to determine the elastic modulus:

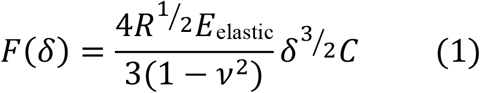

**F** is the applied force, **δ** is the indentation, **R** is the relative radius of the spherical probe (Equation 2), **ν** is the Poisson’s ratio (assumed to be 0.5 for incompressible materials), **C** is a thin-layer correction factor relating indentation depth, tip radius, and sample thickness [33]. **R** accounts for the curvature of the probe tip and cell at the point of contact based on **h** (the height of the cell) [31]:

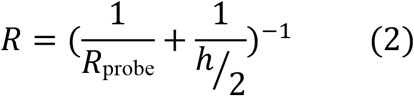

The cell height was significantly decreased after fMLP treatment, but no difference was found between WT and Vcl^-/-^ neutrophils.

### Neutrophil sequestration in mice

The Lifespan Animal Welfare Committee approved all animal studies. Mice were housed in a specific pathogen-free facility at Rhode Island Hospital. Mice harboring floxed *Vcl* alleles (Vcl^f/f^) were kindly provided by Dr. Robert Ross (UC-San Diego) as previously described [34]. Vcl^f/f^ mice were crossed with Mx1-Cre (Mx1^cre^) mice (The Jackson Laboratory) in which Cre recombinase expression is controlled by the Mx1 promoter and can be induced by interferon production after administration of synthetic double-stranded RNA (PolyI: C) [35]. To generate mixed chimeric mice harboring multiple neutrophil genotypes, 8-to 12-week-old C57BL/6 mice (The Jackson Laboratory) were lethally irradiated (10 Gy, single dose) and then reconstituted by intravenous injection of bone marrow cells from a Vcl^f/f^Mx1^cre^GFP^+^ mouse expressing enhanced green fluorescent protein (EGFP) under the ubiquitin C promoter (The Jackson Laboratory) and a Vcl^f/f^ (GFP^-^) control mouse at 1:1 ratio. Disrupting the *Vcl* gene encoding vinculin was induced by intraperitoneal injection of 250 µg of polyinosinic–polycytidylic acid (InvivoGen), three doses, each two days apart, starting four weeks after irradiation, inducing near-complete loss of the respective protein in neutrophils [29]. Murine mixed chimeras were administered 5 µg/kg GM-CSF intravenously while anesthetized using a cocktail of ketamine (125 mg/kg) and xylazine (12.5 mg/kg). Peripheral blood was sampled via saphenous venipuncture. The inferior vena cava was excised to allow pulmonary circulation to flow out from the heart, and the lungs were perfused of unbound cells with 5 mL of HBSS^++^ via the right ventricle. Lungs were harvested and briefly rinsed in HBSS^++^ before digestion according to the Lung Digestion Kit manufacturer’s protocol (Miltenyi). We analyzed the frequency of WT (GFP^-^) and Vcl^-/-^ (GFP^+^) neutrophils in blood and lung tissue samples using flow cytometry.

### Phagocytosis assay

Differentiated (day five) neutrophils were washed 3X in PBS. 2 × 10^6^ neutrophils were resuspended in a 100 μl mixture of pHrodo Green S. *aureus* bioparticles (0.3 mg/mL), FBS (10% v/v) and HBSS^++^. The samples were incubated for 0 or 90 min at 37°C. To quench neutrophil phagocytosis, 1 mL of ice-cold PEB (PBS, 1 mM EDTA, 1% BSA) was added to the samples and placed on ice. To confirm actin-dependent bioparticle internalization, samples were exposed to 10 μg/mL cytochalasin D. After phagocytosis, samples were stained with Zombie Violet and APC-Ly6G and then analyzed by flow cytometry using a MACSQuant 10 (Miltenyi). Data were analyzed to quantify the fraction of Ly6G^high^ neutrophils with a positive pHrodo FITC signal. This experiment was performed with the conditions in duplicate and parallel and repeated across three independent experiments.

### Microfluidic assay

#### Microfluidic device construction

AutoCAD was used to design the microfluidic constriction platforms. Photolithography was performed by Front Range Photomask. The silicon master (SU-8) was then manufactured at the Microfabrication Core Facility at Harvard University. Silanization of the silicon wafer was performed to passivate the surface and allow for easy release of the PDMS. To silanize the master, 20 μL of the silanizing agent — trichloro(1H, 1H, 2H, 2H-perfluorooctyl)silane — was placed next to the silicon wafer within a vacuum desiccator for 30 min. Vacuum was applied for 30 min to form the silane monolayer on the master surface. Finally, the silicon wafer was placed on a hotplate at 150°C for 10 min to evaporate the excess silane.

To prepare the PDMS, a 10:1 ratio of the elastomer to curing agent was mixed and then subjected to degas for an hour. The PDMS was then poured over the silicon master and then baked overnight at 60°C on a hotplate to harden the polymer. The PDMS stamps with the molded microfluidic design imprinted on them were cut out and holes were punched using a blunt 18G needle for inlet and outlet tubing. The surface of the PDMS was cleaned with Scotch tape. Glass coverslips are soaked in concentrated sulfuric acid overnight, rinsed with copious amounts of water (18 Ω), and then rinsed with 70% ethanol solution and dried. The PDMS stamp (design side up) and the glass coverslip were exposed to the corona discharge generated by a Tesla coil (Model BD-20, Electro-Technic Products, Inc.) for two minutes. The PDMS stamp and the glass coverslip were then sandwiched together (design facing the glass) to form the whole microfluidic device. The microfluidic platforms were left overnight to ensure strong adherence.

#### Neutrophil transit through constrictions

A microfluidic device with constrictions 5 μm across and 8 μm high was designed for this assay, adapted from a design kindly provided by Dr. Amy Rowat. The transit of neutrophils through the constrictions was imaged using fluorescence microscopy. For schematic A of Figure 1, hydrostatic pressure was used to drive the neutrophils through the device. A buffer column set at 10 cmH_2_O was used and the fluid velocity at the constrictions was measured to be 0.026 mm/s. Images were acquired at a frame rate of 10 frames/s for 20 min. When experiments were performed using the microfluidic device shown in schematic B of Figure 1, a syringe pump was used due to the high fluid resistance resulting from the mixing channel. The flow rate set on the syringe pump (Braintree Scientific) was 500 nl/min for two syringes for a total of 1 μL/min throughout the device. This resulted in an average fluid velocity of 0.116 mm/s, which is similar to the reported values for fluid velocity in the pulmonary capillaries, ranging from 0.11 mm/s – 0.28 mm/s[15, 36, 37]. Images were acquired at a frame rate of 19 frames/s for 20 min for these experiments. These data were exported as TIFF files to ImageJ, where the tracks were built with the plugin Trackmate [38, 39].

**Figure 1.**
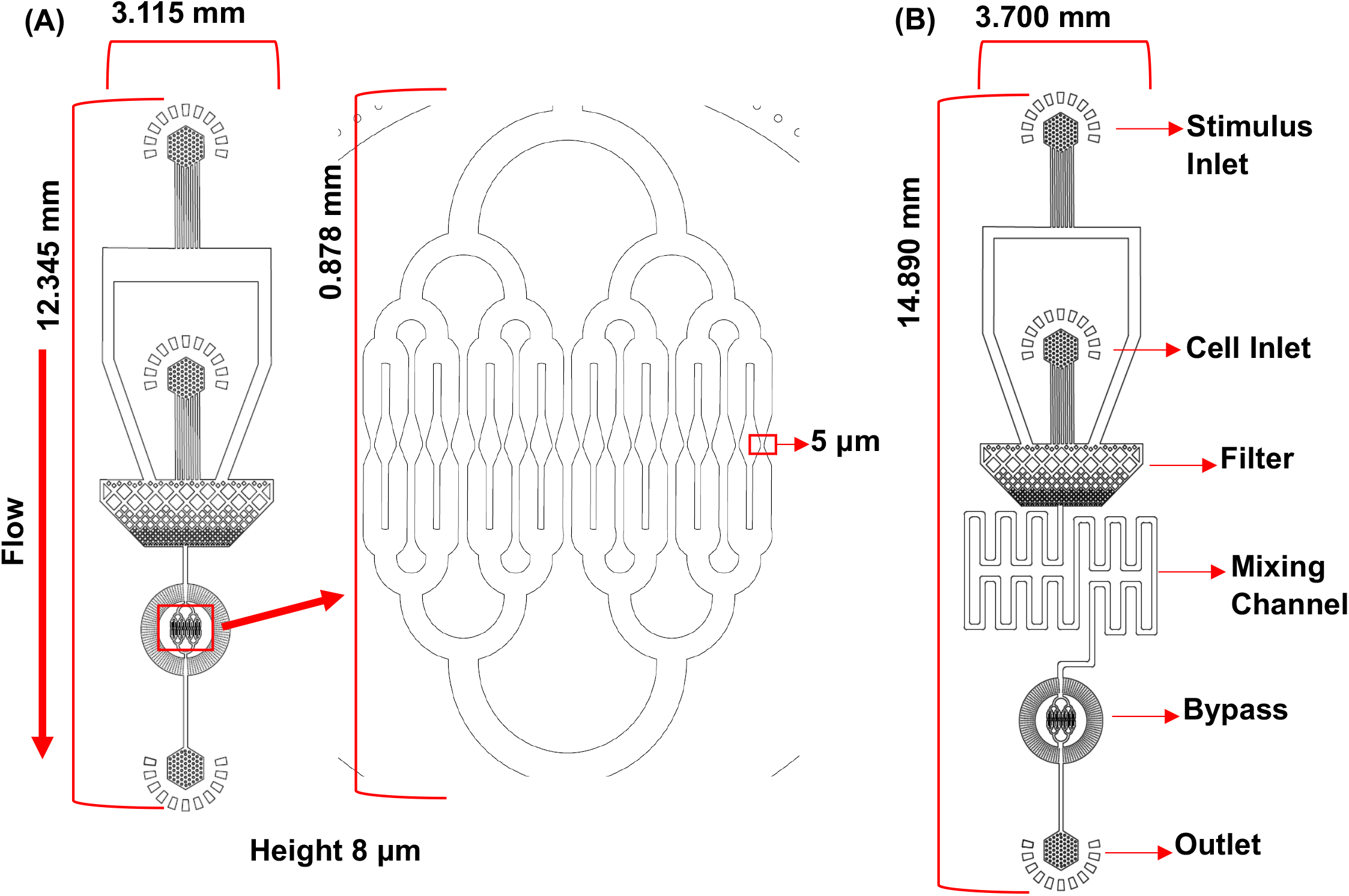
Schematics of microfluidic device designs. (A) Design and layout of the microfluidic device used in the bone marrow neutrophil experiments. The driving pressure was 10 cmH_2_O. (B) Microfluidic design used for studies using HoxB8-conditional progenitor-derived neutrophils. In this design a mixing channel is included to ensure uniform stimulus exposure. A syringe pump set to 500 nl/min was used to perfuse the device at a constant flow rate.

#### Data analysis - constriction microfluidics

The Fiji plugin TrackMate [38] was used for analysis to identify and locate the neutrophils and link them into tracks over time. Each track was visually inspected to confirm the linkages. Tracks were then transferred into Mathematica for final filtering. This filtering is based on a series of gates that the neutrophil must pass to be counted. The first gate is 50 pixels from the array entrance, the second gate is 20 pixels before the center of the constriction, and the final gate is 50 pixels from the end of the array.

Additionally, data were filtered according to cell diameter. The estimated diameters calculated by TrackMate are based on finding the edge of the cell by contrast [38]. We select a wide range of 4 to 12 microns, which results in +/-4 microns around the estimated median. There are 0.437 microns/pixel. Calculating the neutrophil’s diameter is susceptible to pixels turning on and off, and we find it is only a rough estimate of the cell’s actual size. Flow cytometry, in which we can identify mature neutrophils via labeling, is a more accurate estimate for neutrophil size, as shown in Supplemental Figure 1. The result of the filtering described above leaves us with a series of tracks where we can determine the transit time of each neutrophil as a correlation to the cell stifness.

Neutrophil transit times from three independent experiments were pooled. Using Excel, we randomly selected 185 transit times from the pooled data. We use the following macro: **=INDEX($A$1:$A$530**,**RANK(B1**,**$B$1:$B$530)**. We also assign a random distribution of real numbers 0 to 1, using the **RAND()** function, to our pooled data set to prevent duplicates. This method ensures that the data is evenly weighted across our comparisons. These data were then exported back into Mathematica, where their cumulative distributions were plotted against time to yield final kinetic curves. The final fits and statistical analysis were performed with Prism.

#### Neutrophil preparation

On the day of the microfluidic experiment, differentiated neutrophils were washed 3X in optimal buffer (PBS, 1 mM EDTA, 2% FBS). Neutrophils were then purified using an EasySep Mouse Neutrophil Enrichment Kit (StemCell Technologies). The cell samples were always kept on ice until used in the microfluidic device. 1×10^6^ cells were removed from the purified neutrophil mixture, washed 2X with PBS, and resuspended in 200 μL of PBS. These cells were stained with CFSE for 15 min, washed 2X with PBS, and counted. They were then resuspended in filtered (0.22 μm) HBSS^++^, 0.1% Pluronic solution, 1% BSA, and 3 μg/mL DNAaseI so that the final concentration was equal to 1.0×10^6^ cells/mL. In a separate tube, the stimulus mixture was prepared with HBSS^++^, 0.1% Pluronic solution, 1% BSA, 3 μg/mL DNAaseI and 2 μM fMLP. The stimulus of the neutrophils is achieved within the microfluidic device by having two inlets — one for the neutrophils and the other for the stimulus. The final concentration of the stimulus on-chip was 1 μM. All solutions were at room temperature except the optimal buffer, which was ice-cold. The neutrophil and stimulus solutions were drawn into 1 mL syringes with 26G needles. The syringes were attached to the microfluidic chip with polyethylene (PE 50) tubing.

#### Microscopy

The samples were imaged with a TILL Photonics iMIC microscope (FEI Company) with an Olympus 10X UPlanFI objective, NA 0.30. The camera was an Andor iXon3 EMCCD camera. We capture an image every 0.053 s for 20 min.

### Statistical analyses

To determine which model represents the data accurately, the 95% confidence bands were plotted around the fits. Next, the residuals of the fit were inspected to check for a random distribution across the x-axis (all residuals for the fitted curves are in Figure S1). Finally, a calculation was performed of the Extra sum-of-squares F test with the rule for selection of the simpler model unless the P-value is greater than 0.05. To compare the best fits between two different models, an unpaired, parametric t-test was performed and the standard deviation of the residuals (Sy.x) was used to estimate goodness-of-fit. Three independent experiments were performed and all experimental conditions were evaluated in parallel within each experiment. For bone marrow neutrophil studies, neutrophils were purified from three different mice.

Neutrophil phagocytosis experiments were analyzed using a 2-way ANOVA with a Dunnet multiple comparisons test.

## Results

### Design of a microfluidic cell constriction device

To probe neutrophil transit through constrictions that model the dimensions of pulmonary capillaries, we utilized two different microfluidic devices adapted from Nguyen and Hoelzle [40, 41] (Figure 1). The initial design, Figure 1A, has a stimulus inlet, a cell inlet, a filter region, a flow bypass ring, and the single-cell constriction array. The design in Figure 1B has the same features and also includes a long mixing channel to ensure adequate and consistent stimulus exposure for the neutrophils. The filter region prevents debris from the inlets from traveling further into the microfluidic device and also disperses clumps of neutrophils so that only single-cell species traverse through the constriction array. Details about the experiments using each of the device designs are provided below with the support of neutrophil transit data. The narrowest widths in the constriction array are 5 μm and the height across the device is 8 μm.

Figure 2A shows an example of a CFSE-stained neutrophil transiting a constriction (Video 1). The constriction array through which the neutrophils pass is imaged for 20 min, and then image sequences are subjected to analysis (Figure 2B, 2C) [39]. The transit times of neutrophils are plotted as cumulative distributions (Figure 3A) and fit to one of two continuous functions. More traditional forms of mathematical description, such as reporting the median or average single-cell transit time do not adequately account for the full range of observed cell transit behavior and therefore result in the loss of the experimental resolution provided by this microfluidic device. The functions are a one-phase exponential rise and a two-phase exponential rise. The mathematical model of the one-phase exponential rise is:

**Figure 2.**
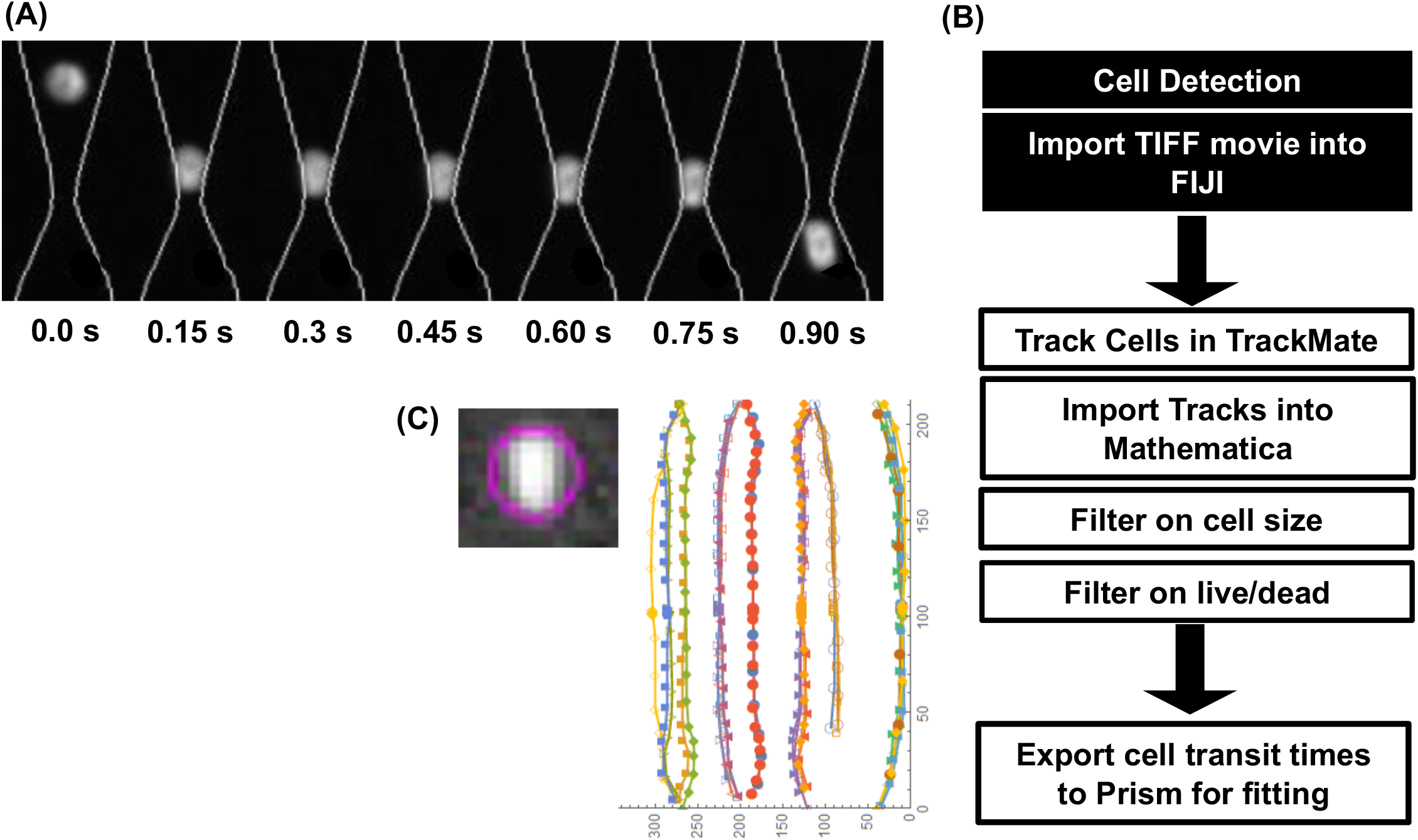
Neutrophil transit data analysis workflow. (A) An example time series of a neutrophil transiting a constriction. (B) The data analysis workflow that yields transit time for each neutrophil passage event that can be exported to Prism for fitting. (C) An example cell position plot from the TrackMate plugin that shows cell identification (highlighted with a purple circle) and the linked tracks for quantifying cell transit over time.

**Figure 3.**
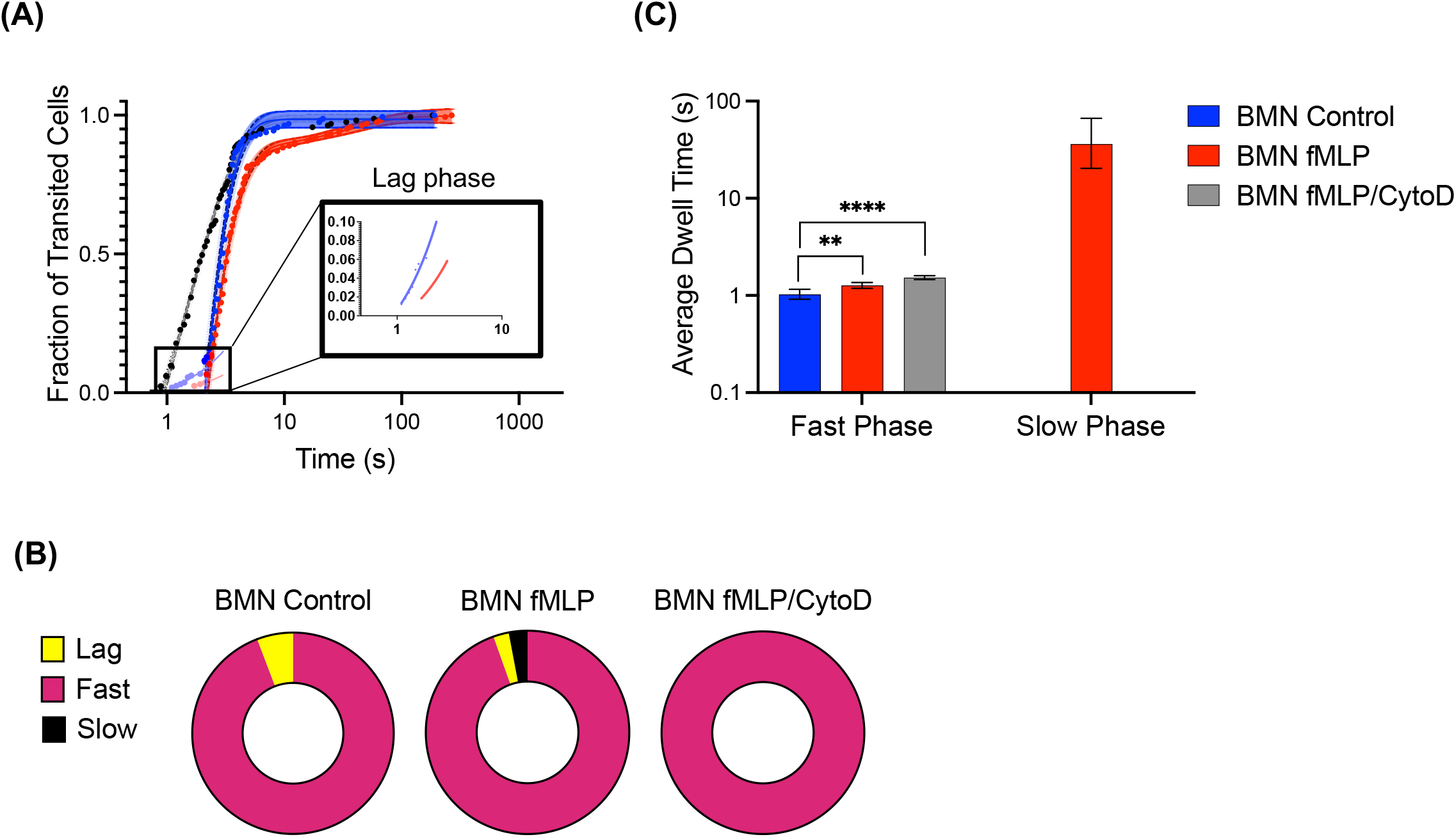
Bone marrow neutrophil transit time distributions and impact of fMLP. (A) Cumulative distributions of BMN transit times for control (blue), stimulated with fMLP (red), and treated with CytoD and fMLP (black). The control group was fit with a linear trend (inset, light blue) and a single-phase exponential rise (dark blue). The fMLP-stimulated group was fit with a linear trend (inset, light red) and a 2-phase exponential rise (dark red). The BMN group treated with CytoD and fMLP was fit with a single-phase exponential rise. (B) The best fit values of the average transit times determined from the fits in (A). The lag phase is associated with the linear fit, the fast phase of the control and CytoD/fMLP data sets is associated with a single exponential rise, and the fast and slow phases for the fMLP-stimulated group data set is associated with a 2-phase exponential rise. (C) The best fit values of the magnitude of the fast phase population determined from the 2-phase exponential rise fits in (A) for the data set of fMLP-stimulated BMNs. Since the control and the CytoD/fMLP data sets fit to a single exponential rise, their population was entered as 100% for clarity. (D) The population distribution of each group, recalculated to include the lag phase proportion of the control and fMLP-stimulated group data sets.

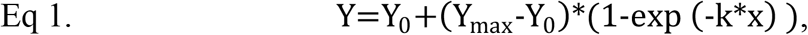

and a two-phase association,

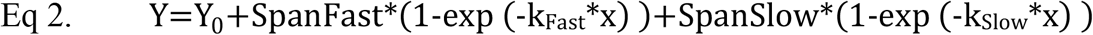

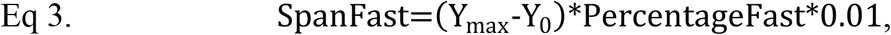

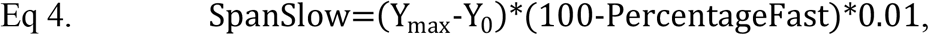

Where each of the terms represents the following: ***Y*** is cell fraction, ***Y***_***0***_ is the cell fraction at 0s, ***Y***_***max***_ is the plateau (maximum cell fraction), ***k*** is the rate constant and its reciprocal, 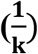, is the average transit time, and ***x*** is time (s). We use the Extra sum-of-squares F-test to choose the mathematical model that best describes the response to each condition. By fitting the neutrophil transit time population trends, we can determine if there are multiple neutrophil populations within each condition and their associated mean transit times.

### Bone marrow neutrophil transit through model capillary segments

We measured the transit times of murine bone marrow neutrophils (BMNs) using the microfluidic platform shown in Figure 1A. Samples were exposed to either control conditions (blue curve) or stimulated with 1 μM fMLP (red curve). Neutrophils were treated with cytochalasin D (CytoD) in combination with fMLP to demonstrate the dependence of neutrophil stiffening on F-actin polymerization (black curve). CytoD is a membrane-permeable fungal toxin that binds to the barbed end of actin filaments, inhibiting both the association and dissociation of subunits.

Neutrophil transit time curves under control and fMLP stimulus conditions exhibit a lag phase, representing 6% and 3% of the neutrophil population, respectively (Figure 3A inset and 3B). The lag phase describes the fastest transiting events under each condition and is likely due to the underlying resistance to deformation under the flow and pressure conditions used. This results in neutrophils transiting at the times represented in the lag phase being rare. The linear slopes between these two conditions are not significantly different, with a p-value of 0.14. This suggests that the deformability characteristics of these rare, fast transiting neutrophils under these two conditions are similar. After the lag phase, rates accelerate and the transit time curve is best described with an exponential rise function.

For the control condition, the acceleration of the population through the constrictions is fully described by a single-phase exponential rise (Eq. 1, Figure 3A). The average transit time for this one-phase exponential is 1.0 s [95% CI 0.92 – 1.2] (Figure 3C). Thus, the control neutrophils have two populations characterized by a lag phase and an exponential rise. The transit time curve for neutrophils exposed to CytoD and fMLP fits a single exponential phase with an average transit time of 1.5 s [95% CI 1.4 – 1.6] (Figure 3C).

For neutrophils stimulated with fMLP, the transit time curve has three phases: a lag and two exponential rise phases that were designated as “slow” and “fast”. The two-phase exponential has 97% [95% CI 97 – 98] of the neutrophil population in the fast phase with an average transit time of 1.3 s [95% CI 1.2 – 1.4] and 2.5% of the population in the slow phase with an average transit time of 36 s [95% CI 20 – 67] (Figure 3B, 3C).

Together, our analyses indicate that the transit of BMNs through narrow constrictions can be described by fitting their distribution curves to a continuous function with multiple phases. By quantifying neutrophil transit times under various conditions, we identify three distinct subpopulations of neutrophils. Our data indicates that neutrophils predominantly transit through a model capillary unit with characteristic “fast” kinetics, regardless of the presence of fMLP as an activating stimulus. A minority of neutrophils exhibits transit kinetics characterized by a lag phase that is abolished by disrupting the actin cytoskeleton. Finally, stimulation of neutrophils with fMLP induces a small population of neutrophils with slow transit. This experimental approach provides a platform for probing the mechanisms by which neutrophils modulate their deformability.

### Transit behavior of neutrophils derived from progenitor cell lines

To further investigate the mechanisms of actin polymerization and remodeling and regulation of neutrophil stifness, we derived neutrophils from HoxB8-conditional progenitors. This experimental system allows for the generation of knockout cell lines that can be maintained in their progenitor state and subsequently differentiated into mature neutrophils. First, we used our microfluidic platform and analyses to establish that neutrophils derived from HoxB8-conditional progenitors transit through constrictions and respond to stimuli similarly to BMNs.

In experiments with BMNs, we observed that fewer than 5% of neutrophils exhibited fMLP-induced activation features, such as the neutrophil transit with slow phase kinetics. In subsequent experiments, we aimed to more deliberately expose cells to the stimulus within the microfluidics prior to reaching the constriction channel. Microfluidic design B includes a long mixing channel to ensure adequate exposure to the stimulus (Figure 1). In the supplemental data, we show the distribution of fluorescein before and after the mixing channel (Figure S2), indicating thorough mixing and a uniform exposure to stimulus in this device. Due to the extremely small dimensions of the microfluidic device, the addition of this channel added significant flow resistance to the experiment. We therefore used a syringe pump in experiments employing microfluidic design B to overcome this resistance and instead set the flow rate to match the fluid flow velocity to that of the capillaries in the lung, estimated between 109 μm/s – 280 μm/s [15, 36, 37]. We performed calibration assays with fluorescent beads and neutrophils to confirm the flow velocity of 130 μm/s through the constriction array for the systemic 1μL/min flow rate (Figure S3).

We perfused wild-type (WT) neutrophils differentiated from HoxB8-conditional progenitors through the microfluidic platform and characterized their transit time distribution. In analyses of the distribution of neutrophil transit times, a single exponential curve describes the transit times of unstimulated neutrophils, while fMLP-stimulated neutrophils require two a two-phase exponential rise (Figure 4A, 4B). Neutrophils treated with CytoD exhibited transit times characterized by a single-phase exponential curve. In contrast to BMNs, these studies yielded transit time distributions that did not exhibit a lag phase. This may be due to the change in microfluidic design and implementation. The average transit time for control neutrophils and neutrophils exposed to fMLP and CytoD are 1.0 s [95% CI 0.98 – 1.1] and 0.21 s [95% CI 0.17 – 0.26], respectively (Figure 4C). For neutrophils stimulated with fMLP, the average transit time within the fast phase is 0.46 s [95% CI 0.41 – 0.52], while the slow phase had an average transit time of 5.2 s [95% CI 4.5 – 6.1] (Figure 4C). Though the transit times are shorter than those for BMNs as expected due to the increased flow rates, only 67% [95% CI 64 – 69] of the fMLP-stimulated neutrophils populate the fast phase (Figure 4B).

**Figure 4.**
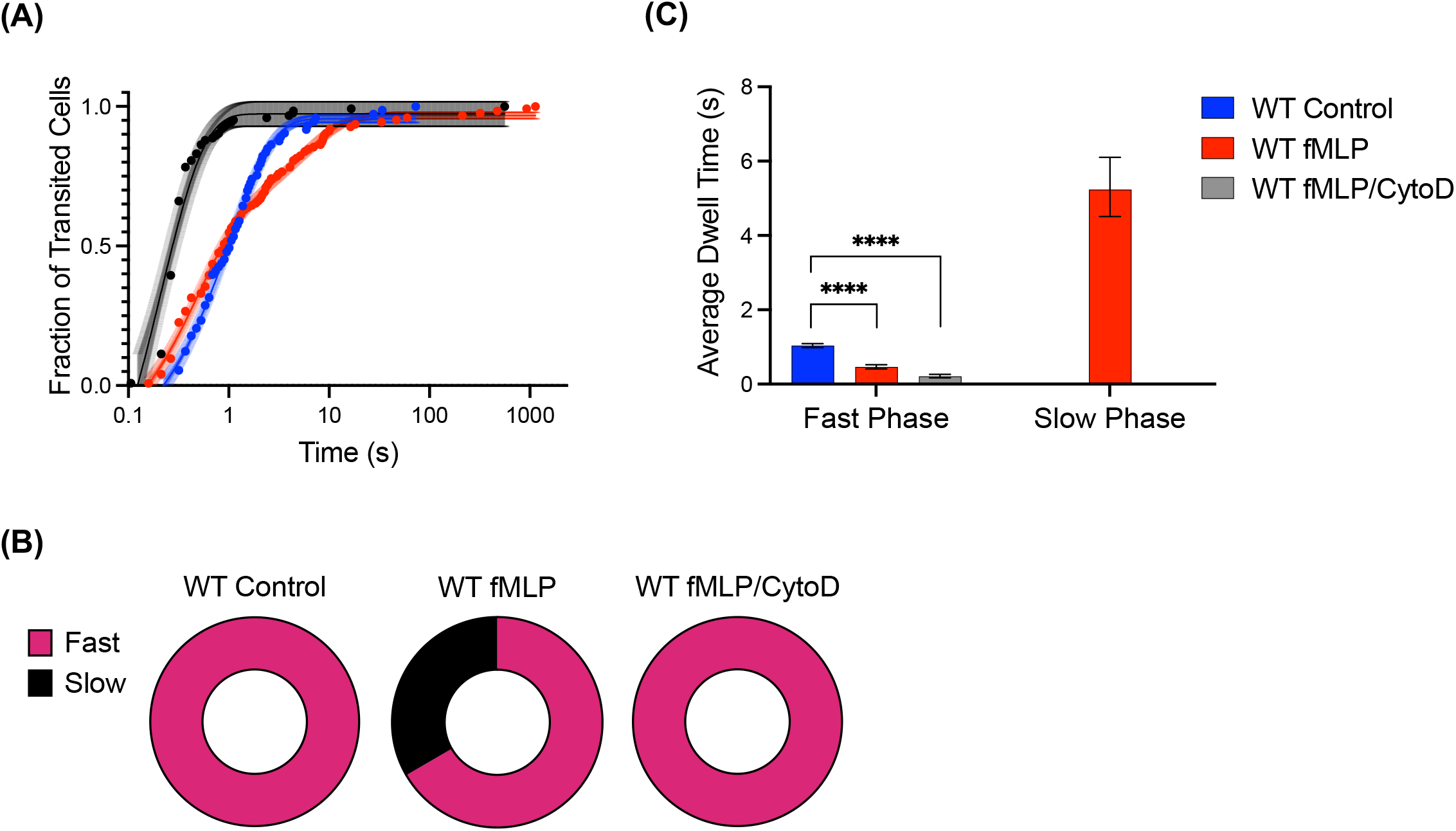
Transit time distributions of wild-type (WT) neutrophils derived from HoxB8-conditional progenitors. (A) The cumulative distributions of transit times for WT control (blue), WT stimulated with fMLP (red), and WT treated with CytoD and fMLP (black). The WT control group was fit with a single-phase exponential rise. The fMLP-stimulated group was fit with a 2-phase exponential rise. The CytoD/fMLP group was fit with a single-phase exponential rise. (B) The best fit values of the average transit times determined from the fits in (A). (C) The population distributions of WT neutrophils under each condition, determined from the best fit values in (A).

### Vinculin regulates neutrophil deformability and transit through 5-μm constrictions

Vinculin is an actin-binding protein whose role in cortical actin organization is well understood in mesenchymal cells but less so in neutrophils [28, 29]. Using CRISPR/Cas9 to generate Vcl^-/-^ progenitors which were then differentiated into neutrophils, we investigated the role of vinculin in neutrophil stiffening and transit through model capillaries.

We perfused WT and Vcl^-/-^ neutrophils through the microfluidic platform and recorded the transit times (Figure 5A). Analyses of transit time curves indicates that WT neutrophils stimulated with fMLP were the only group that had a two-phase transit time distribution (Figure 5B), as described above. In contrast, Vcl^-/-^ neutrophils stimulated with fMLP have a single phase with an average transit time of 0.65 s [95% CI 0.60 – 0.70] (Figure 5C). Unstimulated Vcl^-/-^ neutrophils have an average transit time of 0.57 s [95% CI 0.51 – 0.64] (Figure 5C). While the average dwell time within the fast phase of fMLP-stimulated WT neutrophils is less than that of Vcl^-/-^ neutrophils, because all Vcl^-/-^ neutrophil transit events exist within the fast phase regime their overall transit is faster than that of WT neutrophils. Finally, we observed the shortest transit times for Vcl^-/-^ neutrophils pretreated with CytoD and stimulated with fMLP, with an average dwell time of 0.12 s [95% CI 0.11 – 0.13] (Figure 5C). These data indicate that vinculin plays a crucial role in inducing slow-transiting neutrophils in response to fMLP.

**Figure 5.**
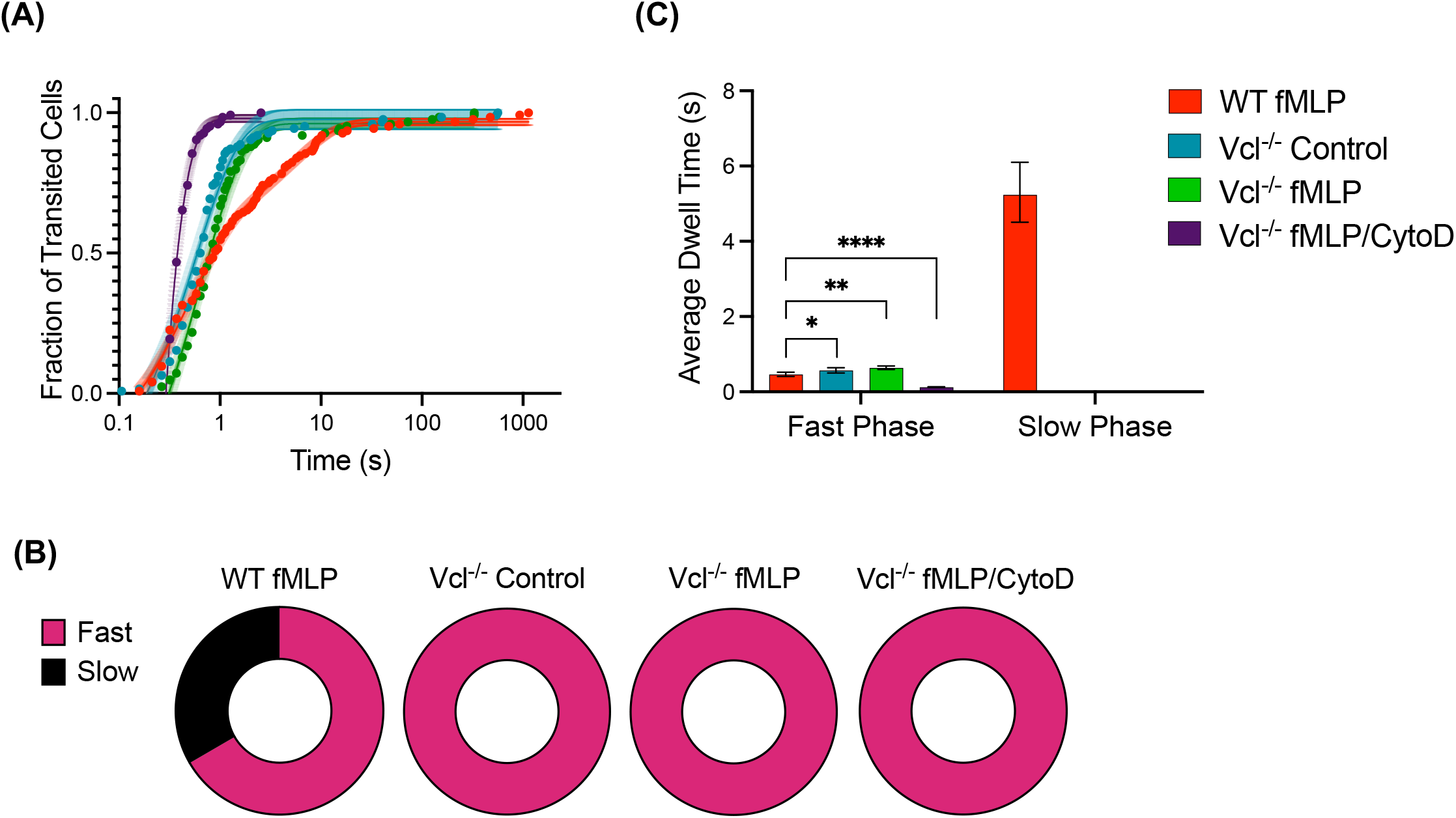
Transit time distributions of WT and Vcl^-/-^ neutrophils. (A) The cumulative distributions of transit times for WT stimulated with fMLP (red), Vcl^-/-^ stimulated with fMLP (green), Vcl^-/-^ control (aqua), and Vcl^-/-^ treated with CytoD and fMLP (purple). The group of WT neutrophils treated with fMLP was fit with a 2-phase exponential rise. All other data sets in this figure were fully described by a single-phase exponential rise. (B) Comparison among groups of the best fit values of the average transit times determined from the fits in (A).

The development of resistance to changes in deformability prolong neutrophil transit in the pulmonary microvasculature. To directly probe neutrophil deformability, we performed analyses of neutrophils derived from HoxB8-conditional progenitors using atomic force microscopy. In these experiments, neutrophils were allowed to settle on a glass coverslip and remained in a rounded morphology, as indicated by a constant cell height across all experimental conditions (Figure 6A). In the absence of stimulus, WT and Vcl^-/-^ neutrophils had a similar elastic modulus (Figure 6B). We found that Vcl^-/-^ neutrophils have a lower elastic modulus than WT neutrophils when stimulated with fMLP (Figure 6B). Together, these data indicate that the faster transit properties of Vcl^-/-^ neutrophils are likely due to an abrogated stiffening response to activation-induced actin remodeling.

**Figure 6.**
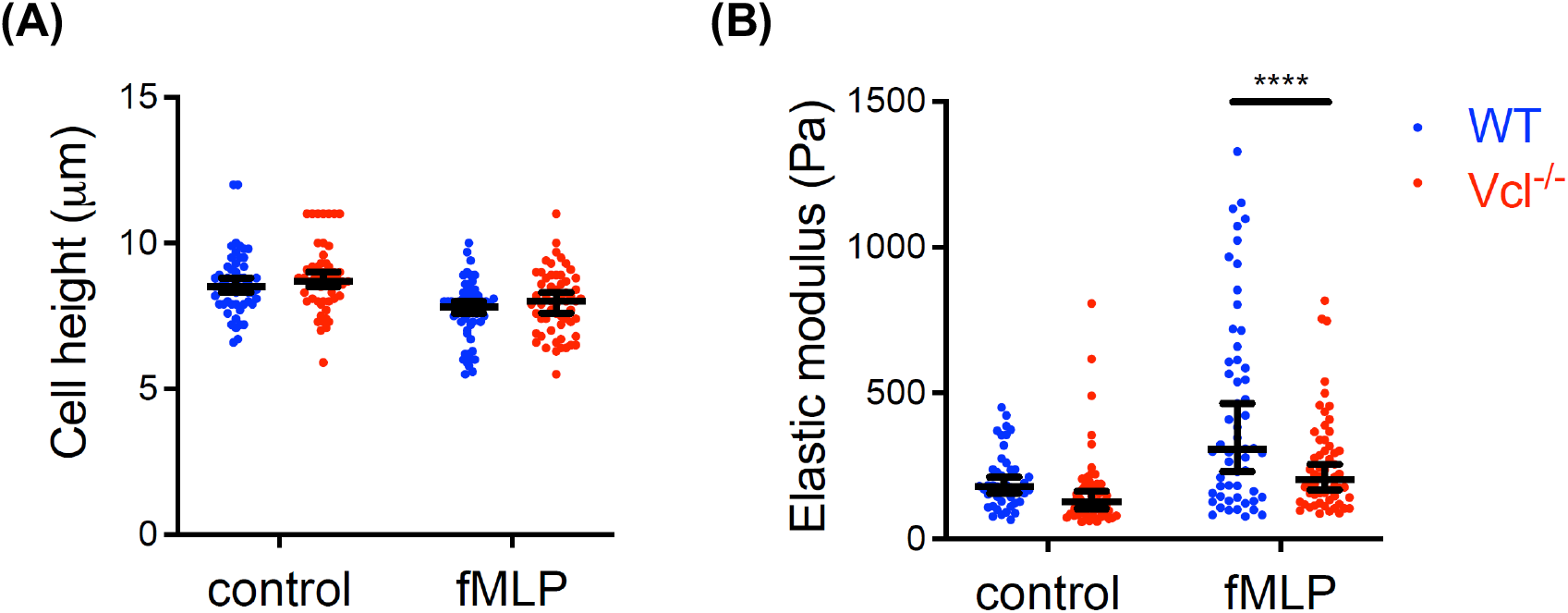
AFM analyses of WT and Vcl^-/-^ neutrophils. Neutrophils were allowed to settle on coverslips and then exposed to control conditions or 1 μM fMLP. (A) The cell height and (B) elastic modulus (Pa) for each cell was measured by AFM as described in the Methods. Measurements from each cell are single data points on the plots shown. n > 56 cells per group. Data were analyzed using two-way ANOVA with Tukey multiple comparisons test. ****p < 0.001.

### Vinculin and neutrophil sequestration in the lungs

To evaluate the impact of vinculin-dependent neutrophil stiffening on in the acute accumulation of neutrophils in the lungs, we performed studies in an *in vivo* mouse model of pulmonary neutrophil sequestration. Experiments utilized mixed chimeric mice that harbor both WT and Vcl^-/-^ neutrophils, distinguished by the co-expression of GFP [29]. We administered granulocyte-macrophage cell-stimulating factor (GM-CSF) intravenously to induce neutrophil priming, stiffening, and sequestration [42]. After evacuating the lungs of non-sequestered neutrophils, we analyzed the remaining WT and Vcl^-/-^ neutrophils and compared their relative frequencies to those observed in the circulation (Figure 7). These data, expressed as the frequency of Vcl^-/-^ neutrophils relative to WT, indicate that GM-CSF induces the sequestration of more WT neutrophils relative to Vcl^-/-^ neutrophils in the lungs, as compared to their ratio in the blood (Figure 7). These data demonstrate that vinculin plays a role in the acute GM-CSF-induced sequestration of neutrophils in the lung, and are consistent with the phenotype of vinculin-deficient neutrophils lacking a defined population with slow phase transit kinetics in microfluidic analyses.

**Figure 7.**
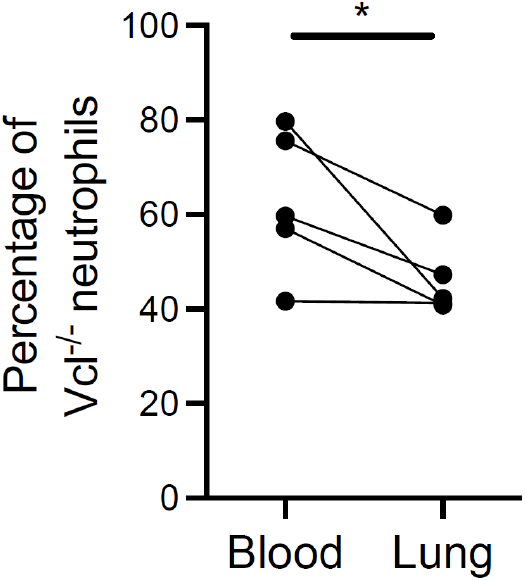
GM-CSF-induced neutrophil sequestration in the lungs of mixed chimeric mice. Mice harboring both WT and Vcl^-/-^ neutrophils received intravenous injection of GM-CSF. After 15 min, mice were euthanized and the blood and lungs harvested for analyses. Data are expressed as the percentage of Vcl^-/-^ neutrophils in the blood and lung. n = 5 mice across two independent experiments. Data were analyzed using a paired Student’s t-test. *p < 0.05.

### Vinculin is not essential for neutrophil phagocytosis

Our results indicate that vinculin has a role in the actin-dependent modulation of neutrophil stiffening and transit through narrow constrictions. Given the important role of actin reorganization in other neutrophil functions, we investigated the potential role of vinculin in neutrophil phagocytosis. Phagocytosis is a specialized F-actin-dependent form of endocytosis that is critical for the antimicrobial function of many immune cells, including monocytes, macrophages, neutrophils, dendritic cells, osteoclasts, and eosinophils [43]. To determine whether vinculin plays a role in neutrophil phagocytosis, we performed experiments to evaluate the uptake of non-viable *Staphylococus aureus*. We incubated neutrophils with pHrodo-conjugated *S. aureus* bioparticles and assayed their uptake after 90 min. We found that Vcl^-/-^ neutrophils phagocytosed the microbe just as well as WT neutrophils (Figure 8). As expected, disruption of the actin cytoskeleton with CytoD abrogated phagocytosis (Figure 8). Thus, while neutrophil stiffening in response to fMLP is inhibited by the loss of vinculin, actin-dependent phagocytosis is maintained.

**Figure 8.**
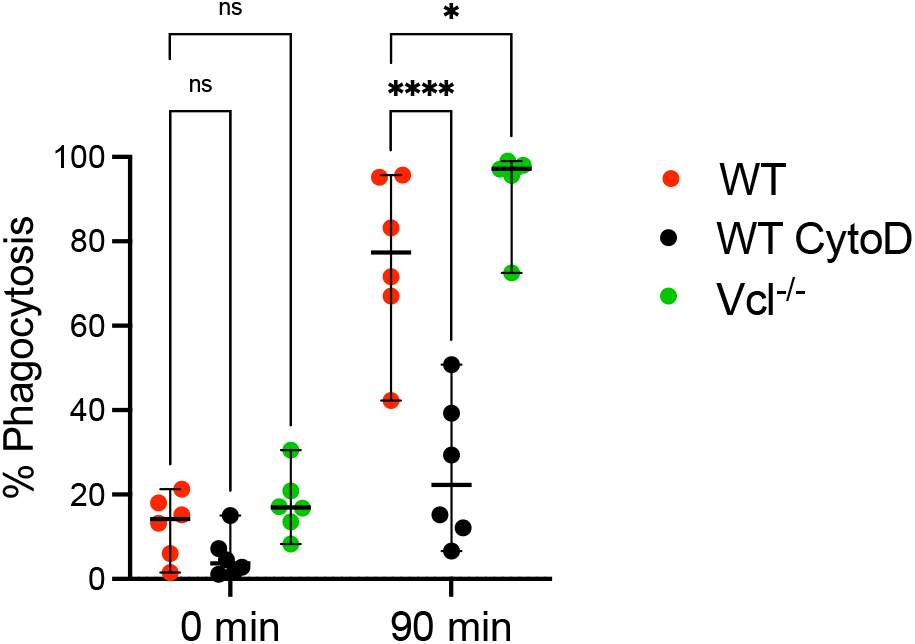
Neutrophil phagocytosis of *S. aureus* bioparticles. WT and Vcl^-/-^ neutrophils derived from HoxB8-conditional progenitors were incubated with pHrodo Green-conjugated *S. aureus* bioparticles for 90 min at 37°C and analyzed by flow cytometry to quantify the percentage of viable neutrophils that successfully internalized *S. aureus*. To demonstrate the dependence of phagocytosis on the actin cytoskeleton, a group of samples were incubated with 10 µg/mL CytoD. For the 0 min time point, the reaction is immediately quenched after addition of *S. aureus*.

## Discussion

The extensive microvasculature of the lungs provides the massive surface area necessary for blood-gas exchange to oxygenate tissues throughout the body. The circulation is also the highway by which immune cells survey and traffic to tissue sites. Neutrophils, as the most abundant leukocyte in the circulation, frequently pass through the pulmonary capillary network and must deform to do so. As we observe in our microfluidic model of capillary segments, the process of neutrophil passage slows their transit relative to the bulk flow, a phenomenon that results in the enrichment of neutrophils within the pulmonary capillaries relative to other vascular beds, even at homeostasis when neutrophils are uniformly in a basal, inactive state [10]. Under conditions in which neutrophils receive activating signals, either from local inflammation of the lung or a systemic inflammatory response, neutrophils accumulate in the pulmonary capillaries [10, 23]. It was subsequently established that physical trapping of neutrophils in the lung capillaries is an actin-dependent process and leads to their retention [10, 23]. Specifically, physical trapping due to cortical actin assembly-induced neutrophil stiffening mediates initial sequestration, whereas β2 integrins play a role in prolonging neutrophil retention [26]. While these broader mechanisms have been known for almost 30 years, the specific intracellular mediators of actin reorganization that drive neutrophil stiffening still remain unclear.

Our studies aimed to take advantage of microfluidics to precisely design features at scale and thereby establish a tractable experimental system to probe the mechanisms of neutrophil transit in model pulmonary capillary units. Constriction microfluidics enables quantification of the functional elastic modulus, or stiffness, of the whole body of the cell by applying the pressure of fluid perfusion to force it to deform in order to passage through a narrow channel. An advantage of the design flexibility of microfluidics is that by using a separate inlet for the stimulus, one can control the timing for exposure of neutrophils to a stimulus and ensure that each cell experiences the chemical for a similar amount of time prior to encountering the constriction channel. Thus, these experiments allow a neutrophil to respond to both a biochemical signal and mechanical deformation via the microfluidic device. This model, which mimics the pulmonary capillary system, is not achievable with the classical methods of testing mechanical properties such as AFM and cell aspiration [44]. Additionally, this approach allows for the examination of many more events than what is feasible with other techniques. The microfluidic mid-throughput assay allows us to collect single-cell mechanical data on hundreds to thousands of cells, yielding robust data collection that then provides access to more sophisticated data analysis techniques in which one can analyze population-based effects.

In our study, the number of neutrophil transit events analyzed in each data set allowed us to confidently fit the data to continuous functions. By taking advantage of data analysis techniques optimized with single-particle tracking, we fit the cumulative distribution curves of transit times to exponential rise functions. Under certain experimental conditions, it was necessary to employ more than one exponential rise to accurately fit the data, suggesting that distinct populations of neutrophils give rise to that particular trend. This is illustrated by the need for at least a 2-phase exponential function to fit the data for BMN transit in the presence of fMLP stimulus. Analyses of BMN transit time indicated that the addition of fMLP results in a subset of neutrophils undergoing stiffening that acted to increase their transit time 28-fold compared to neutrophils that did not respond. It is possible that there may be additional biologically distinct populations than we resolve using this approach, but many more neutrophil transit events would need to be analyzed to gain the statistical power to distinguish them.

In our analyses of BMNs, the transit time curves of both the control and fMLP-stimulated group have a lag phase. By determining the x-intercept of the two lag phases of the control and stimulus and the x-intercept of the exponential rise of the negative control, we observe that a little less than a second is the fastest any cell can transit under these conditions. In addition, the lag phase of the stimulus and control groups indicates that the basal neutrophil stifness results in a shift that increases the minimum transit time compared to the negative control group that was exposed to CytoD to disrupt the actin cytoskeleton. In other words, the basal tone of neutrophils results in a bottleneck that makes it rare to observe a cell transit faster than 1.5 s, which is in contrast to neutrophils treated with CytoD in which over 30% of the neutrophils have transited in less than 1.5 s.

Similar to the neutrophil transit subpopulation dynamics observed in studies with BMNs, analyses of neutrophils derived from HoxB8-conditional progenitors revealed the fMLP-induced appearance of a stiffened neutrophil subset whose transit times were characterized by a second exponential phase of the curve fit. By introducing a stimulus-mixing channel into the microfluidic platform, we found an increased proportion of neutrophils in the subset described by slow phase transit kinetics in response to fMLP. It is intriguing that despite uniform exposure to the stimulus, only a fraction of neutrophils exhibit slower transit kinetics. This was not a feature unique to the microfluidic system, as AFM analyses also suggested that only a proportion of neutrophils shift towards a drastically increased elastic modulus in response to fMLP. Together, these findings indicate the flexibility and robustness of using microfluidics to probe neutrophil passage through narrow channels, and also establish that neutrophils derived from HoxB8-conditional progenitors modulate their deformability in response to stimuli in a similar manner as primary neutrophils.

In previous studies, our lab established that vinculin plays a role in neutrophil adhesion and motility that depends upon the experimental context (e.g., presence of fluid flow) [29]. While those studies probed integrin-dependent neutrophil motility on two-dimensional surfaces, here we focused on the potential role of vinculin in aspects of neutrophil trafficking that occur while a neutrophil is still in a round morphology, as does while circulating. We found that vinculin deficiency abrogated the neutrophil stiffening response to fMLP, manifested by Vcl^-/-^ neutrophils not developing a second population with slow phase transit kinetics in response to fMLP. Instead, the average transit time of fMLP-stimulated Vcl^-/-^ neutrophils (0.6 s) was similar to that of unstimulated WT (1.0 s) and of unstimulated Vcl^-/-^ neutrophils (0.6 s), differences that are not likely to be biologically meaningful. Interestingly, vinculin-deficient neutrophils whose actin cytoskeleton was disrupted with CytoD exhibited the shortest average transit times of all conditions evaluated. This suggests that Vcl^-/-^ neutrophils retain some basal level of actin-dependent stifness that may be organized in structures that do not involve vinculin.

Neutrophil stiffening and sequestration in pulmonary capillaries represents a physiological mechanism for host defense of the lungs [10]. However, under pathophysiologic conditions such as those present in ARDS, continued accumulation of neutrophils in the pulmonary capillary bed and their premature deployment of antimicrobial defenses contributes to injury of host tissues [5, 9]. Here, we demonstrate that vinculin plays a role in neutrophil sequestration *in vivo* in response to GM-CSF, an agent that rapidly primes neutrophils and induces their transient retention in the lung [42]. Understanding the mechanisms of pathophysiological neutrophil sequestration in the lungs, and whether they are distinct from those that we identify in this study, should be the subject of future investigation. We observed that Vcl^-/-^ neutrophils are able to engulf and internalize *S. aureus*, indicating that the actin-dependent process of phagocytosis that is key for intracellular eradication of microbes does not require vinculin. This suggests the feasibility of therapeutically targeting neutrophil stiffening while sparing important host defense functions of neutrophils.

## Acknowledgments

We thank Dr. Paul Ekert, Dr. Patrice Dubreuil, and Dr. Amy Rowat for kindly providing reagents, resources, or advice to conduct the studies.

## Funding

This research was supported by awards from the National Institutes of Health:

R35GM124911 to C.T.L.; T32HL134625 supported B.M.N., R25HL088992 supported Y.R. and K.A..

## Author Contributions

B.M.N.: Conceiving and designing experiments, conducting experiments, analyzing data, writing and editing the manuscript.

Z.S.W.: Conceiving and designing experiments, conducting experiments, analyzing data.

K.A.: Conducting experiments.

Y.R.: Conducting experiments.

M.K.S.: Helping to conduct and analyze AFM experiments.

E.M.D.: Supervising AFM experiments.

C.T.L.: Supervising the project, conceiving and designing experiments, writing and editing the manuscript.

## References

1. Eworuke, E., Major, J. M., McClain, L. I. G. National incidence rates for Acute Respiratory Distress Syndrome (ARDS) and ARDS cause-specific factors in the United States (2006-2014). Journal of Critical Care. 2018;47, 192–197.

2. Bellani, G., Laffey, J. G., Pham, T., et al. Epidemiology, Patterns of Care, and Mortality for Patients With Acute Respiratory Distress Syndrome in Intensive Care Units in 50 Countries. Jama-Journal of the American Medical Association. 2016;315, 788–800.

3. Biehl, M., Ahmed, A., Kashyap, R., et al. The Incremental Burden of Acute Respiratory Distress Syndrome: Long-term Follow-up of a Population-Based Nested Case-Control Study. Mayo Clin Proc. 2018;93, 445–452.

4. Alipanah, N. and Calfee, C. S. Phenotyping in acute respiratory distress syndrome: state of the art and clinical implications. Current Opinion in Critical Care. 2022;28, 1–8.

5. Yang, S. C., Tsai, Y. F., Pan, Y. L., et al. Understanding the role of neutrophils in acute respiratory distress syndrome. Biomedical Journal. 2021;44, 439–446.

6. Aulakh, G. K. Neutrophils in the lung: “the first responders”. Cell Tissue Res. 2018;371, 577–588.

7. Palakshappa, J. A., Krall, J. T. W., Belfield, L. T., et al. Long-Term Outcomes in Acute Respiratory Distress Syndrome Epidemiology, Mechanisms, and Patient Evaluation. Critical Care Clinics. 2021;37, 895–911.

8. Chiumello, D., Coppola, S., Froio, S., et al. What’s Next After ARDS: Long-Term Outcomes. Respiratory Care. 2016;61, 689.

9. Grommes, J. and Soehnlein, O. Contribution of neutrophils to acute lung injury. Mol Med. 2011;17, 293–307.

10. Doerschuk, C. M. Mechanisms of leukocyte sequestration in inflamed lungs. Microcirculation. 2001;8, 71–88.

11. Doerschuk, C. M. Neutrophil rheology and transit through capillaries and sinusoids. Am J Respir Crit Care Med. 1999;159, 1693–5.

12. Reutershan, J. and Ley, K. Bench-to-bedside review: Acute respiratory distress syndrome how neutrophils migrate into the lung. Critical Care. 2004;8, 453–461.

13. Shirai, A. Modeling neutrophil transport in pulmonary capillaries. Respiratory Physiology & Neurobiology. 2008;163, 158–165.

14. Ekpenyong, A. E., Toepfner, N., Fiddler, C., et al. Mechanical deformation induces depolarization of neutrophils. Sci Adv. 2017;3, e1602536.

15. Ekpenyong, A. E., Toepfner, N., Chilvers, E. R., et al. Mechanotransduction in neutrophil activation and deactivation. Biochim Biophys Acta. 2015;1853, 3105–16.

16. Looney, M. R. and Bhattacharya, J. (2014) Live Imaging of the Lung. In Annual Review of Physiology, Vol 76, Volume 76 (D. Julius, ed) 431–445.

17. Mizgerd, J. P., Meek, B. B., Kutkoski, G. J., et al. Selectins and neutrophil traffic: Margination and Streptococcus pneumoniae-induced emigration in murine lungs. Journal of Experimental Medicine. 1996;184, 639–645.

18. Bullard, D. C., Kunkel, E. J., Kubo, H., et al. Infections and deficiency of leukocyte rolling and recruitment in E-/P-selectin mutant mice. Faseb Journal. 1996;10, 1605–1605.

19. Doerschuk, C. M., Winn, R. K., Coxson, H. O., et al. CD18-DEPENDENT AND CD18-INDEPENDENT MECHANISMS OF NEUTROPHIL EMIGRATION IN THE PULMONARY AND SYSTEMIC MICROCIRCULATION OF RABBITS. Journal of Immunology. 1990;144, 2327–2333.

20. Diz-Munoz, A., Fletcher, D. A., Weiner, O. D. Use the force: membrane tension as an organizer of cell shape and motility. Trends Cell Biol. 2013;23, 47–53.

21. Marucha, P., Mercado, A., Fernandez, M. REGULATION OF ACTIN EXPRESSION IN GM-CSF-INDUCED PMN. Journal of Dental Research. 1995;74, 130–130.

22. Bochsler, P. N., Neilsen, N. R., Dean, D. F., et al. STIMULUS-DEPENDENT ACTIN POLYMERIZATION IN BOVINE NEUTROPHILS. Inflammation. 1992;16, 383–392.

23. Worthen, G. S., Schwab, B., Elson, E. L., et al. MECHANICS OF STIMULATED NEUTROPHILS - CELL STIFFENING INDUCES RETENTION IN CAPILLARIES. Science. 1989;245, 183–186.

24. Gebb, S. A., Graham, J. A., Hanger, C. C., et al. SITES OF LEUKOCYTE SEQUESTRATION IN THE PULMONARY MICROCIRCULATION. Journal of Applied Physiology. 1995;79, 493–497.

25. Andonegui, G., Bonder, C. S., Green, F., et al. Endothelium-derived Toll-like receptor-4 is the key molecule in LPS-induced neutrophil sequestration into lungs. J Clin Invest. 2003;111, 1011–20.

26. Doerschuk, C. M. THE ROLE OF CD18-MEDIATED ADHESION IN NEUTROPHIL SEQUESTRATION INDUCED BY INFUSION OF ACTIVATED PLASMA IN RABBITS. American Journal of Respiratory Cell and Molecular Biology. 1992;7, 140–148.

27. Peng, X., Nelson, E. S., Maiers, J. L., et al. New insights into vinculin function and regulation. Int Rev Cell Mol Biol. 2011;287, 191–231.

28. Cohen, J. T., Danise, M., Hinman, K. D., et al. Engraftment, Fate, and Function of HoxB8-Conditional Neutrophil Progenitors in the Unconditioned Murine Host. Frontiers in Cell and Developmental Biology. 2022;10.

29. Wilson, Z. S., Witt, H., Hazlett, L., et al. Context-Dependent Role of Vinculin in Neutrophil Adhesion, Motility and Trafficking. Sci Rep. 2020;10, 2142.

30. Wang, G. G., Calvo, K. R., Pasillas, M. P., et al. Quantitative production of macrophages or neutrophils ex vivo using conditional Hoxb8. Nature Methods. 2006;3, 287–293.

31. Kanthilal, M. and Darling, E. M. Characterization of mechanical and regenerative properties of human, adipose stromal cells. Cell Mol Bioeng. 2014;7, 585–597.

32. Darling, E. M., Zauscher, S., Block, J. A., et al. A thin-layer model for viscoelastic, stress-relaxation testing of cells using atomic force microscopy: do cell properties reflect metastatic potential? Biophys J. 2007;92, 1784–91.

33. Dimitriadis, E. K., Horkay, F., Maresca, J., et al. Determination of elastic moduli of thin layers of soft material using the atomic force microscope. Biophys J. 2002;82, 2798–810.

34. Zemljic-Harpf, A. E., Miller, J. C., Henderson, S. A., et al. Cardiac-myocyte-specific excision of the vinculin gene disrupts cellular junctions, causing sudden death or dilated cardiomyopathy. Mol Cell Biol. 2007;27, 7522–37.

35. Velasco-Hernandez, T., Säwén, P., Bryder, D., et al. Potential Pitfalls of the Mx1-Cre System: Implications for Experimental Modeling of Normal and Malignant Hematopoiesis. Stem cell reports. 2016;7, 11–18.

36. Wagner, W. W., Latham, L. P., Gillespie, M. N., et al. DIRECT MEASUREMENT OF PULMONARY CAPILLARY TRANSIT TIMES. Science. 1982;218, 379–381.

37. Eichhorn, M. E., Ney, L., Massberg, S., et al. Platelet kinetics in the pulmonary microcirculation in vivo assessed by intravital microscopy. Journal of Vascular Research. 2002;39, 330–339.

38. Tinevez, J.-Y., Perry, N., Schindelin, J., et al. TrackMate: An open and extensible platform for single-particle tracking. Methods. 2017;115, 80–90.

39. Schindelin, J., Arganda-Carreras, I., Frise, E., et al. Fiji: an open-source platform for biological-image analysis. Nature Methods. 2012;9, 676–682.

40. Nguyen, A. V., Nyberg, K. D., Scott, M. B., et al. Stiffness of pancreatic cancer cells is associated with increased invasive potential. Integr Biol (Camb). 2016;8, 1232–1245.

41. Hoelzle, D. J., Varghese, B. A., Chan, C. K., et al. A Microfluidic Technique to Probe Cell Deformability. Jove-Journal of Visualized Experiments. 2014.

42. Summers, C., Singh, N. R., White, J. F., et al. Pulmonary retention of primed neutrophils: a novel protective host response, which is impaired in the acute respiratory distress syndrome. Thorax. 2014;69, 623–629.

43. Rosales, C. and Uribe-Querol, E. Phagocytosis: A Fundamental Process in Immunity. BioMed research international. 2017;2017, 9042851–9042851.

44. Ren, C. G., Yuan, Q. Y., Braun, M., et al. Leukocyte Cytoskeleton Polarization Is Initiated by Plasma Membrane Curvature from Cell Attachment. Developmental Cell. 2019;49, 206-+.

