## Supplemental Information for "Vinculin plays a role in neutrophil stiffening and transit through model capillary segments"

### Single Phase Exponential Rise

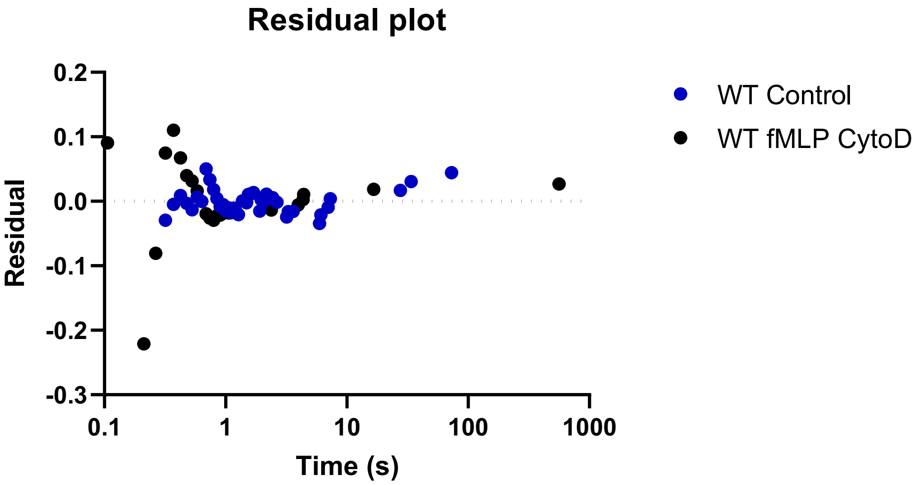


### Two-Phase Exponential Rise

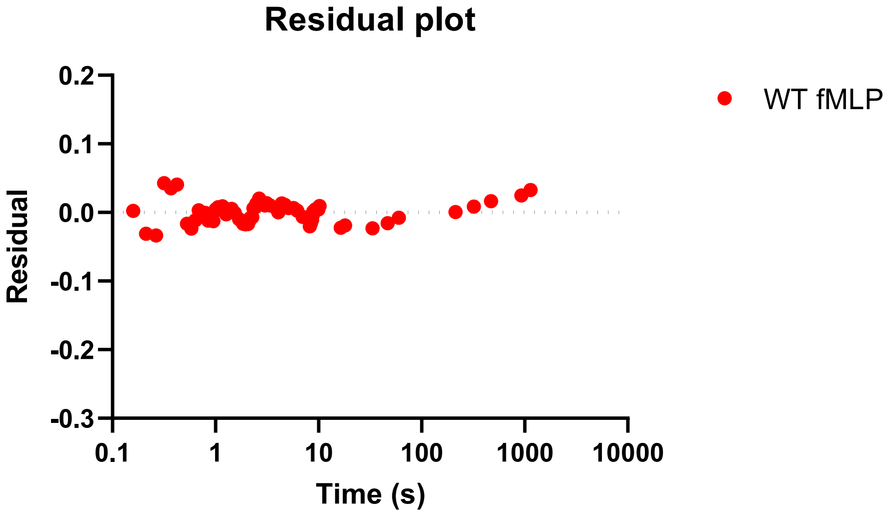


**Figure S1. Residuals of transit time distribution curve fits.** Residuals are plotted and indicate a random distribution across the x-axis.

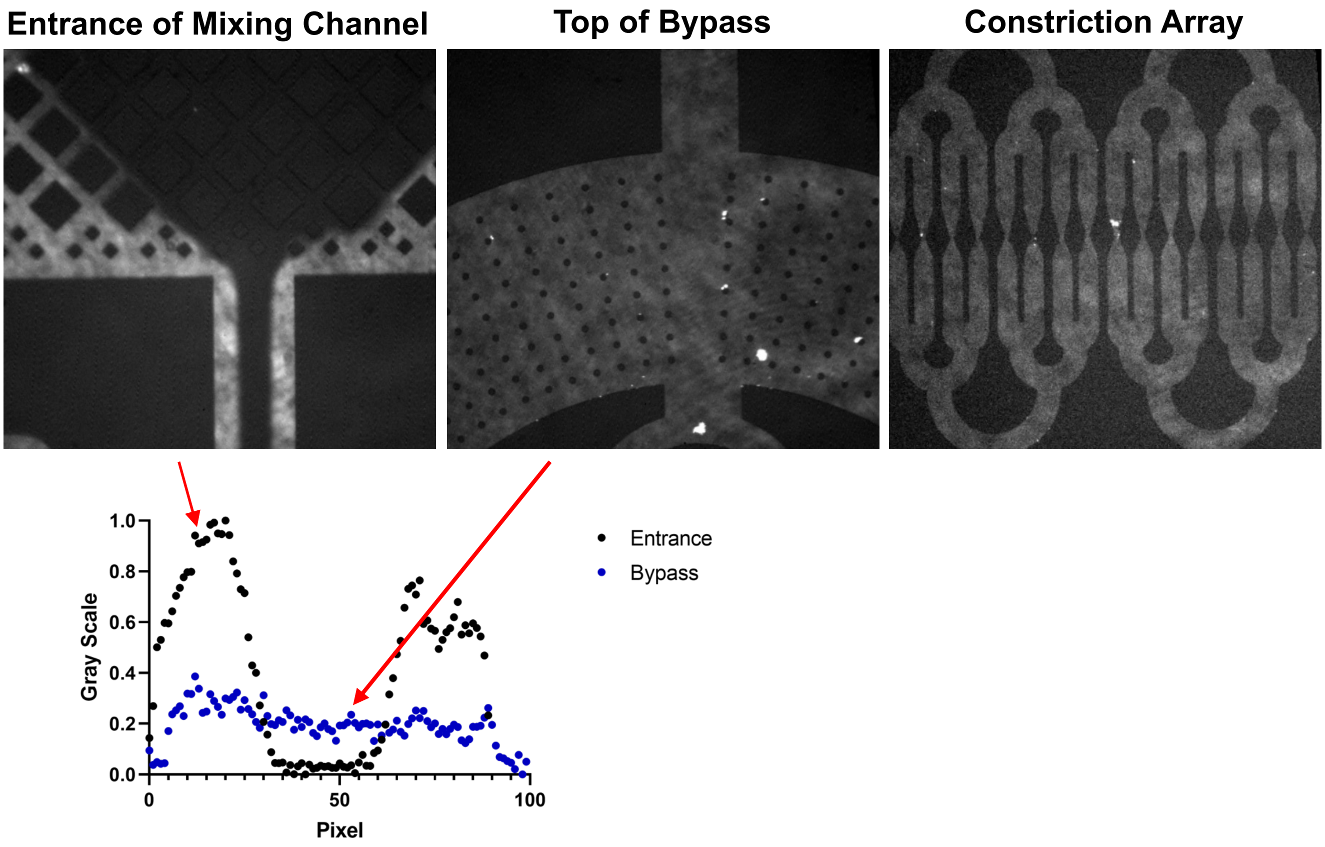


**Figure S2. Microfluidic mixing channel and stimulus distribution.** Fluorescein was perfused through the stimulus inlet under the same conditions used in experiments. The distribution of fluorescein was measured at sections of the microfluidic device before and after the mixing channel.

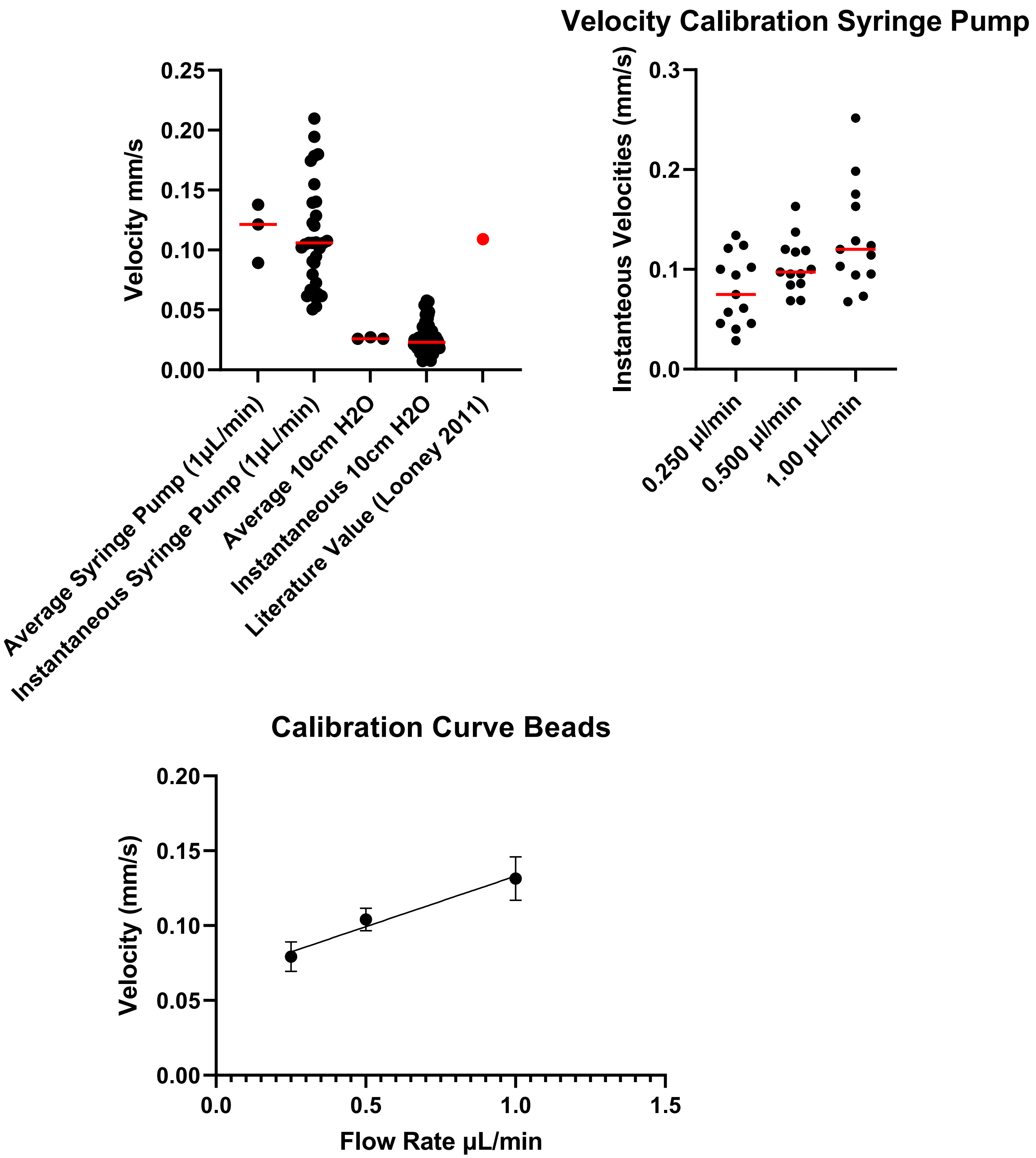


**Figure S3. Calibration of microfluidic perfusion conditions.** Fluorescent beads were perfused through the microfluidic device at the indicated rates (syringe pump) or height of a column of water (constant pressure). The bead velocity was measured by video microscopy.
